# Development of a non-invasive novel individual marmoset holder for evaluation by awake functional magnetic resonance brain imaging

**DOI:** 10.1101/2023.12.21.572749

**Authors:** Fumiko Seki, Terumi Yurimoto, Michiko Kamioka, Takashi Inoue, Yuji Komaki, Atsushi Iriki, Erika Sasaki, Yumiko Yamazaki

**Author notes:** These two authors contributed equally to this work (Fumiko Seki, Terumi Yurimoto). Correspondence authors: Erika Sasaki, Yumiko Yamazaki **Correspondence to:** Erika Sasaki Department of Marmoset Biology and Medicine, Central Institute for Experimental Animals. 3-25-12, Tonomachi, Kawasaki-ku, Kawasaki, 210-0821, Japan.

## Abstract

**Background:** Although functional MRI (fMRI) in awake marmosets (*Callithrix jacchus*) is fascinating for functional brain mapping and evaluation of brain disease models, it is difficult to launch awake fMRI on scanners with less than 15 cm of bore size. A universal marmoset holder for the small-bore size MRI was designed, and evaluated whether this holder could conduct auditory stimulation fMRI in the awake state.

**New Method:** The marmoset holder was designed with an outer diameter of 71.9 mm. A holder was designed to allow adjustment according to the individual head shape, enabling to use the holder universally. An awake fMRI study of auditory response was conducted to evaluate the practicality of the new holder. Whole-brain activation was investigated when marmosets heard the marmoset social communication “phee call,” an artificial tone sound, music (bossa nova), and reversed of those.

**Results:** The prefrontal cortex was significantly activated in response to phee calls, whereas only the auditory cortex was activated in response to pure tones. In response to bossa nova, the marmoset’s visual and auditory cortices were activated. In contrast, the auditory response was decreased when marmosets heard phee calls and bossa nova music played backward. Their stimulus-specific responses indicated they perceived and differentiated sound characteristics in the fMRI environment.

**Comparison with Existing Methods:** A holder does not require surgical intervention or custom-made helmet to minimize head movement in small space.

**Conclusion:** Our newly developed holder made it possible to perform longitudinal fMRI experiments on multiple marmosets in a less invasive manner.

**Highlight:**

- We designed a universal marmoset holder for the optimization of small-bore size MRI. We aimed to evaluate whether the data acquired by fMRI with this holder indicated that awake marmosets were likely to recognize sounds during fMRI and whether the brain activity shown as fMRI signals reflected the characteristics of each sound.
- After the acclimatization of the marmosets to the holder, an awake fMRI study was conducted to evaluate the practicality of the new holder and successfully acquired the auditory responses to the set of sound stimuli.
- Our newly developed holder does not require surgical intervention to minimize head movement, which allows for less invasive fMRI experiments and makes it easier to conduct longitudinal fMRI for multiple marmosets.

## Introduction

Animal models are essential in understanding the mechanisms underlying functional brain activation and intractable human brain diseases (Perovnik et al., 2023). A New World monkey, the common marmoset (*Callithrix jacchus*), is becoming an increasingly attractive animal model because of its practical advantages, such as small body size permitting high-resolution MRI scanning. Additionally, the marmoset’s lissencephaly can easily identify functionally defined areas on the brain’s surface. Furthermore, the topological layout of areas of marmoset cortices closely matches those of humans (Walker et al., 2017), and genetic modification techniques have also been developed to create neuronal disease models using marmosets (Sasaguri et al., 2022; Sato et al., 2016; Sato et al., 2020; Tomioka et al., 2017).

Functional magnetic resonance imaging (fMRI) in marmosets has been used to study sensory systems (Schaeffer et al., 2020a). It has also been used to discover the mechanisms of corticothalamic activation during generalized absence status epilepticus (Tenney et al., 2004) and the neuronal circuits responding to exposure to the psychostimulant drug MDMA (Brevard et al., 2006; Meyer et al., 2006) in a non-invasive manner. Therefore, evaluating the onset and progression of neuronal diseases in marmoset models is useful. One particular advantage of fMRI for neuroimaging is that it allows longitudinal studies to follow age-related changes in animal brain function. Using fMRI in longitudinal studies is crucial because nonhuman primates, unlike mice, have various genetic backgrounds that modify phenotypic responses. fMRI in marmosets has also been improved by the development of optimized hardware, such as the radiofrequency coil (Gilbert et al., 2023; Papoti et al., 2013) and optimized marmoset holders and head-fixation chambers to conduct fully awake MRI (Liu et al., 2013; Schaeffer et al., 2019a). In such environments, a variety of task-based assessments have been established to study visual (Cléry et al., 2020; Hung et al., 2015a, b; Schaeffer et al., 2019b; Schaeffer et al., 2020b), auditory (Jafari et al., 2023; Sadagopan et al., 2015; Toarmino et al., 2017), olfactory (Ferris et al., 2001; Ferris et al., 2004), and tactile processing. Furthermore, brain activity in the resting state has been thoroughly investigated (Belcher et al., 2016; Belcher et al., 2013), and a unique feature of the default mode network in marmosets has been demonstrated (Liu et al., 2019). Functional brain connectivity of open-access resources acquired from fully awake resting-state fMRI data is available (Schaeffer et al., 2022). These are essential databases to understand marmoset brain function and evaluate brain disease models.

However, the number of publications on marmoset fMRI has not increased (Schaeffer et al., 2020a). One possible reason for not using fMRI in marmosets is that it is necessary to have a wide-bore MRI scanner for small animals to replicate currently published methods. In small animal MRI, the bore size in marmosets ranges from 15 to 30 cm without a gradient coil (Schaeffer et al., 2020a). An MRI with a small bore has advantages such as lower costs and a higher potential to equip high gradients than an MRI with a wide bore. Small-bore MRI is also sufficient if the animals are under anesthesia. However, introducing a marmoset holder for fMRI is challenging to apply to small-bore MRI scanners (Schaeffer et al., 2020a). Therefore, if the development of a marmoset holder for small-bore MRI scanners were possible, the opportunities to conduct fMRI in marmosets would expand.

Developing appropriate holding methods for the marmosets to perform fMRI using small-bore MRI scanners is necessary. An important component of marmoset holders is how to fix the head gently yet firmly to prevent head movement because most fMRI studies need to detect brain activity. The major types of head fixation in marmosets are helmet fixation (Silva et al., 2011) and chamber fixation (Schaeffer et al., 2020a). Helmet-type fixation requires the production of customized helmets fabricated using a 3D printer based on individual head morphology data. This method is non-invasive; therefore, longitudinal evaluation over the years is possible. However, a challenge with this type of helmet is that custom-made helmets are difficult to use with multiple individuals, making it labor-intensive to study different marmosets. Chamber-based head fixation applies surgical implementation to the head and does not rely on the shape of the marmoset’s head; therefore, it can be easily applied to any marmoset.

However, this type requires surgery to affix the chamber, which increases the risk of infection and makes long-term evaluation difficult. Accidents involving anesthesia during surgery are also a risk. In particular, genetically modified marmosets, which are precious and rare, should not be lost to these risks.

We designed a universal marmoset holder to optimize small-bore size MRI to address these issues. The parts of the holder were minimized so that the marmosets could be fixed in a narrow space. The holder was designed to allow adjustment according to the individual head shape to fix the head, making it possible to conduct fMRI on all marmosets with a single holder. Additionally, the new method does not require surgery; thus, longitudinal MRI studies over many years are possible.

We aimed to evaluate whether the data acquired by fMRI with this holder indicated that awake marmosets were likely to recognize sounds during fMRI, thus demonstrating that our holder can conduct fMRI in the awake state. If the marmosets could hear sounds during the experiments, the brain activity shown as fMRI signals would reflect the characteristics of each sound. We chose auditory stimuli to evaluate whether the auditory response could be observed even when the acoustic noise produced by the scanner interfered with hearing the sound. We included a social communication call among the various sounds because it concerns the level of the awake state, attention, social experience, and memory, not just passive acoustic perception. A wide range of brain regions can be affected by controlling these conditions, allowing for assessing the detectability and sensitivity of functional images acquired using this method.

## Materials & methods

### Animals

Five healthy adult male marmosets, born and reared at the Central Institute for Experimental Animals (CIEA) or purchased from CLEA Japan, Inc. (Tokyo, Japan), were used in this study (mean age, 77.2 ± 8.6 months; average body weight, 362 ± 15.4 g. The Institutional Animal Care and Use Committee of the CIEA (approval numbers: 20068A, 21129A) approved all experiments.

### Holder design

The animal holder was designed using 3D computer-aided (CAD) software (Fusion 360; Autodesk, San Francisco, CA) to satisfy the following conditions: 1) minimize head movement to < 0.6 mm; 2) maintain the outer diameter at approximately 72 mm to fit within the MRI bore; 3) no invasive surgery required to immobilize the animal; 4) the marmosets can hold a natural resting sphinx position; 5) the vital signs of the animals can be monitored using a medical monitor; and 6) the holder does not interfere with hearing, vision, or olfaction. An MRI-compatible camera (MRC Systems, Heidelberg, Germany), a light source (Machida Endoscope Co., Chiba, Japan), and earphones (STAX SR-003, Stax Ltd., Saitama, Japan) were placed in front of each animal’s face. MRI-compatible earphones were used as speakers. A light source was used to keep the animal awake during the experiment. The earphones were placed 5 cm in front of the marmoset’s face in the bore and adjusted to a volume that could be heard from outside the MRI scanner while running the experiment to provide auditory stimuli (Supplementary Fig. 1). The helmets and U-pins were designed not completely to cover the marmoset’s ears. A hole was made around the ear of the helmet. Therefore, the sound could be heard even when the helmet was on the head.

### Acclimatization to the marmoset holder

All the animals were acclimatized to the experimental conditions described previously before the fMRI experiment (Silva et al., 2011). The marmosets were acclimatized by training once every 2 weeks as follows. The marmoset in the holder was placed in a mock coil, a speaker was placed at the top of the tube, and the camera and light used to record the video were placed in the upper part of the coil. After the room was darkened and the MRI scanner noise from the earphones was played (Supplementary Fig. 2), the training was started. The experimenter evaluated the level of acclimatization for 1–3 days. The scoring criteria were as follows: 1) the marmoset does not move; 2) movement with intermittent struggle; 3) movement with a struggle for approximately half of the acclimatization session. The training duration was 15 minutes. If the experimenter scored the animal with a 1 or 2, the next training session was conducted for 30 min. If the training was performed for 45 min. and the experimenter scored the animal with a 1 or 2, then the fMRI was conducted for the animal. When the animal moved more than 1 voxel during the fMRI experiment (Schaeffer et al., 2020a), the training was repeated for 45 min. We recorded the animals during training using a video camera to confirm that the marmoset was awake. When the marmosets closed their eyelids, we confirmed their movement in response to environmental sounds. A liquid reward was provided after the acclimatization sessions were completed. The experimenter carefully observed the animals before, during, and after acclimatization. To evaluate acclimatization, average estimates of head motion based on fMRI data were compared before and after acclimatization. Head motion was estimated by registering individual fMRI volumes to the middle volume. The body weights of the animals were also measured during training.

### MRI

In this study, a 7.0-T Biospec 70/16 scanner system, equipped with actively shielded gradients at a maximum strength of 700 mT/m, using 72 mm quadrature transmit/receive RF coil (Bruker BioSpin GmbH; Ettlingen, Germany) was used.

### Design of the auditory presentation

The sound sources were novel to the participants and were prepared in three main categories: sounds with high, middle, and low ecological relevance. The sound with high ecological relevance was the marmoset“phee cal” (Miller et al., 2010b), recorded from marmosets living in different breeding rooms from the experimental animals. Bossa nova music (Artists, 2002) was selected for middle ecological relevance because some animals prefer folk music from their original habitat to music from other regions (Watanabe et al., 2018); parts without human vocals were used. Pure-tone sounds with a low ecological relevance were also prepared. The tone frequencies (7 kHz and 1.2 kHz) and duration (1.5 s) were adjusted to make these equivalent to phee calls (Eliades and Tsunada, 2019). In addition, the reverse sounds of the phee calls and bossa nova music were used as auditory stimuli, with the same frequency as the original sounds. All sounds were played repeatedly for less than 20 s at 1 s intervals.

A block design was developed according to a previous report using these sound sources to measure neural responses to auditory stimuli (Toarmino et al., 2017)(Fig. 1). The stimulus was organized into 20-s sound ON and 40-s sound OFF and repeated for 5 min during the fMRI scan. Six runs for each fMRI experiment were performed as follows: 1) phee call; 2) 7 kHz pure tone; 3) bossa nova; 4) phee call played backward; 5) 1.2 kHz pure tone; and 6) bossa nova played backward. An MR-compatible camera and light source were used to monitor the animal during the MRI.

**Figure 1.**
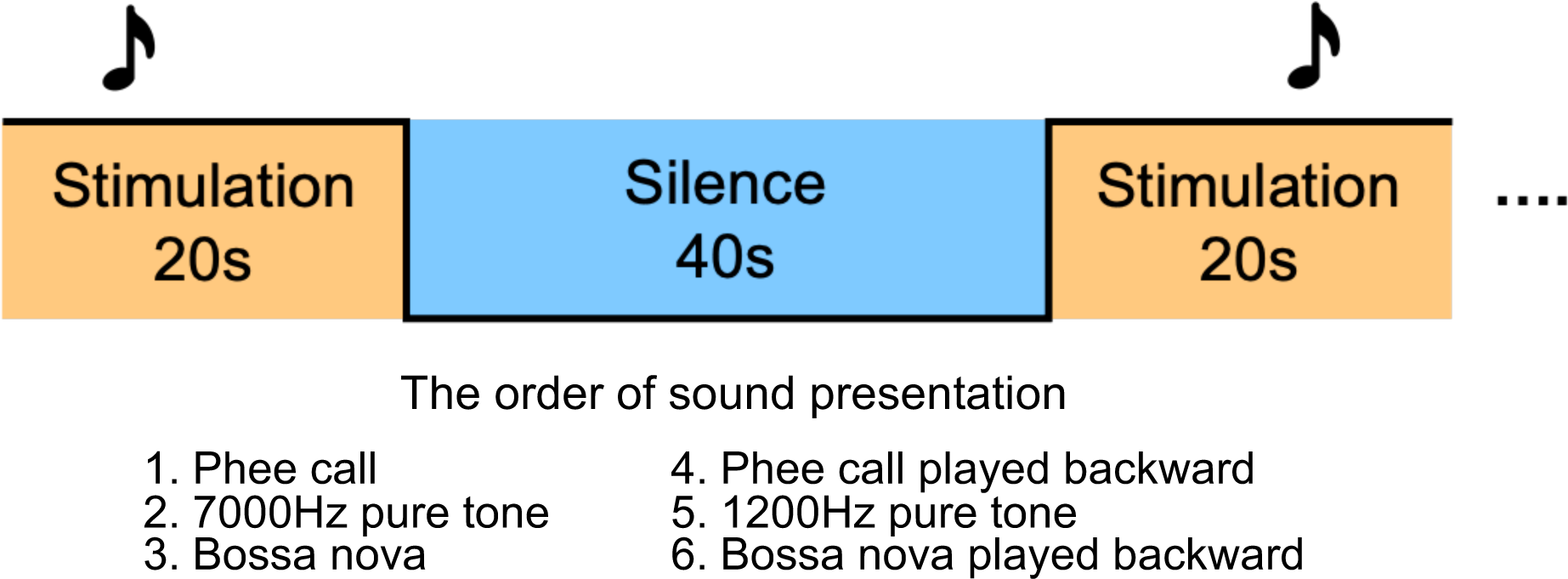
Auditory fMRI block design. Stimulation was presented for 20 s and was silent for 40 s afterward. Six scans were performed for each experiment.

### MRI acquisition and analysis

T2*-weighted functional images were obtained using a gradient-echo echo-planar imaging (EPI) sequence for blood oxygenation level-dependent (BOLD) contrast. The parameters were as follows: repetition time (TR)/echo time (TE): 2,000/15 ms; field of view (FOV): 38.4 × 38.4 mm; 21 axial slices; in-plane image matrix: 64 × 64; spatial resolution: 0. 6 × 0.6 (mm); slice thickness: 1 (mm); flip angle: 60°; repetitions: 150 times; scan time: 5 min. Data with reversed phase-encoding blips, which are images with distortions in the opposite directions, were also collected.

For structural imaging, T2-weighted images (T2WI) were acquired using a rapid acquisition with relaxation enhancement (RARE) sequence with the following parameters: TR/TE: 3,000/30 ms; FOV: 40 × 40 mm; 30 axial slices; image matrix: 160 × 160; spatial resolution: 0. 25 × 0.25 mm; slice thickness: 1 mm; RARE factor, 8; number of averages, 8; scan time, 6 min 30 s.

After correcting for susceptibility distortion (Andersson et al., 2003), the functional images were realigned, slice-time corrected, and registered to the normalized T2WI in the standard space (Hikishima et al., 2011; Seki et al., 2017) using the ANTs pipeline (http://stnava.github. Io/ANTs/). The normalized data were smoothed with an isotropic Gaussian kernel (2.0 mm) and skull-stripped. Functional images were decomposed into spatially independent components using independent component analysis to manually classify and remove noise components (Griffanti et al., 2014). Denoised images were statistically analyzed using FSL FEAT (Woolrich et al., 2001). A one-sample t-test was performed to examine the brain activity induced by the sounds. All statistical images were thresholded with clusters determined by z > 3.1 and a cluster-corrected significance threshold of p < 0.05 (Worsley, 2001). The brain areas significantly activated in response to auditory stimuli were identified using the MRI brain atlas (Hashikawa et al., 2015; Woodward et al., 2018).

## Results

### Development of marmoset holder and acclimatization to the holder

All the hardware parts were created using a 3D printer (Formlabs Inc., Somerville, MA). The holder was designed with an outer diameter of 71.9 mm and a thickness of 3 mm and was made of resin plastic (Fig. 2, Supplementary Fig. 3). The holder consisted of the main holder, helmet, chin rest, collar, back lid, sliding connector, and two U-pins, each fixed with screws or tape. The U-pins restrained the movement of the head, and forward and backward movements of the body were restrained by a collar holding the shoulders. During marmoset restraint, vital status and condition were monitored using a pulse oximeter (Medtronic Japan, Tokyo, Japan), a respiration sensor (SA Instruments, Stony Brook, NY), and a camera. We designed holes in the tube and back lid for this purpose. The helmet was designed to view the face of the marmoset from the front. The back lid also prevented accidental removal of the pulse oximeter from the feet of the animal. The holder parts were fitted to the marmosets as follows. First, the marmoset was passed through an awake tube and restrained using a collar. Second, a helmet was placed on the animal when it was calm. Third, a U-pin was passed through the helmet slide and lightly screwed from the side to restrain head movement completely (Fig. 3). This holder could hold a 480 g marmoset in the sphinx position. All hardware components are available in Supplemental Data 1. The method used to build these parts and fix marmosets to the holder is shown in Supplemental Video 1.

**Figure 2.**
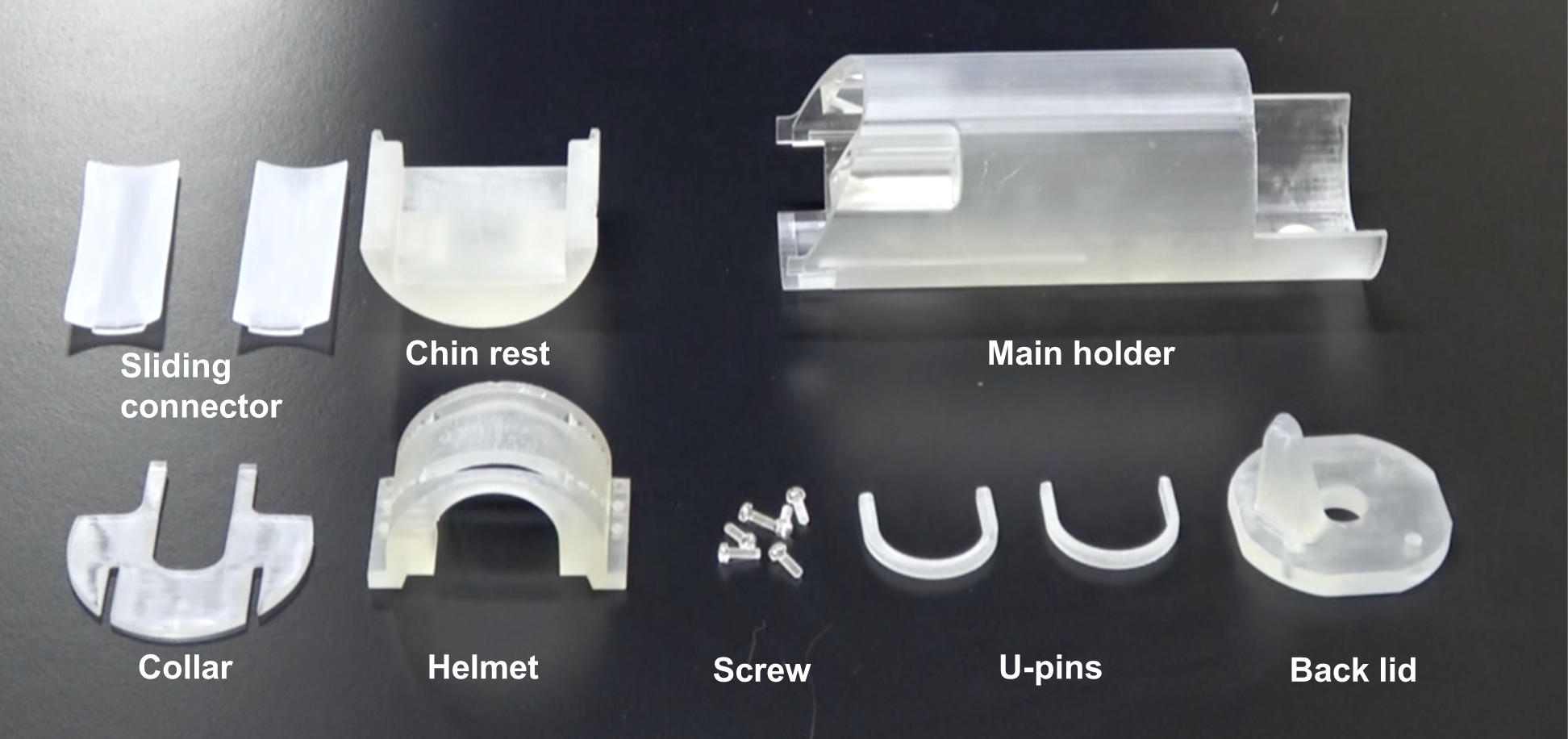
A photograph of the holder parts.

**Figure 3.**
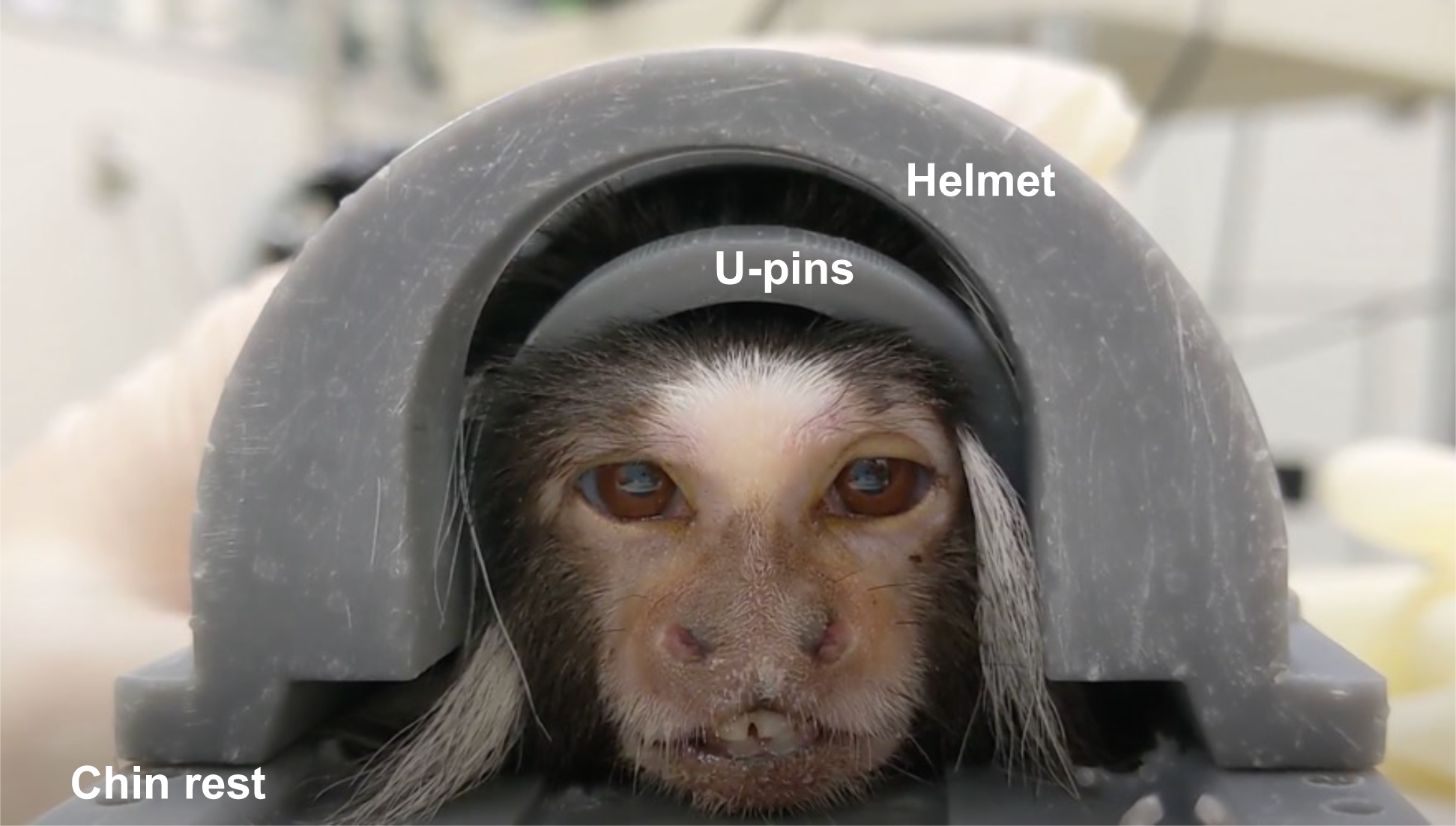
A photograph of a marmoset in the holder.

### Acclimatization to the holder

We confirmed that all marmosets acclimated to the holder. The animals reached the 45 min. session in a mean of 2.4 times. After fully acclimated, the marmosets (except for No. 5) were retrained once every 2 weeks to maintain acclimation until the fMRI experiments started. Although the interval between training sessions varied among the marmosets, each animal was trained approximately 26 times. The body weights of individuals during training before and after the start of MRI acquisition were measured and shown in Fig. 4. The change in body weight during acclimatization was 96–111% compared to that at the start. The average distance of head movements without acclimatization was 0.036 mm in the left-right direction, 0.143 mm in the vertical direction, and 0.22 mm in the front-back direction.

**Figure 4.**
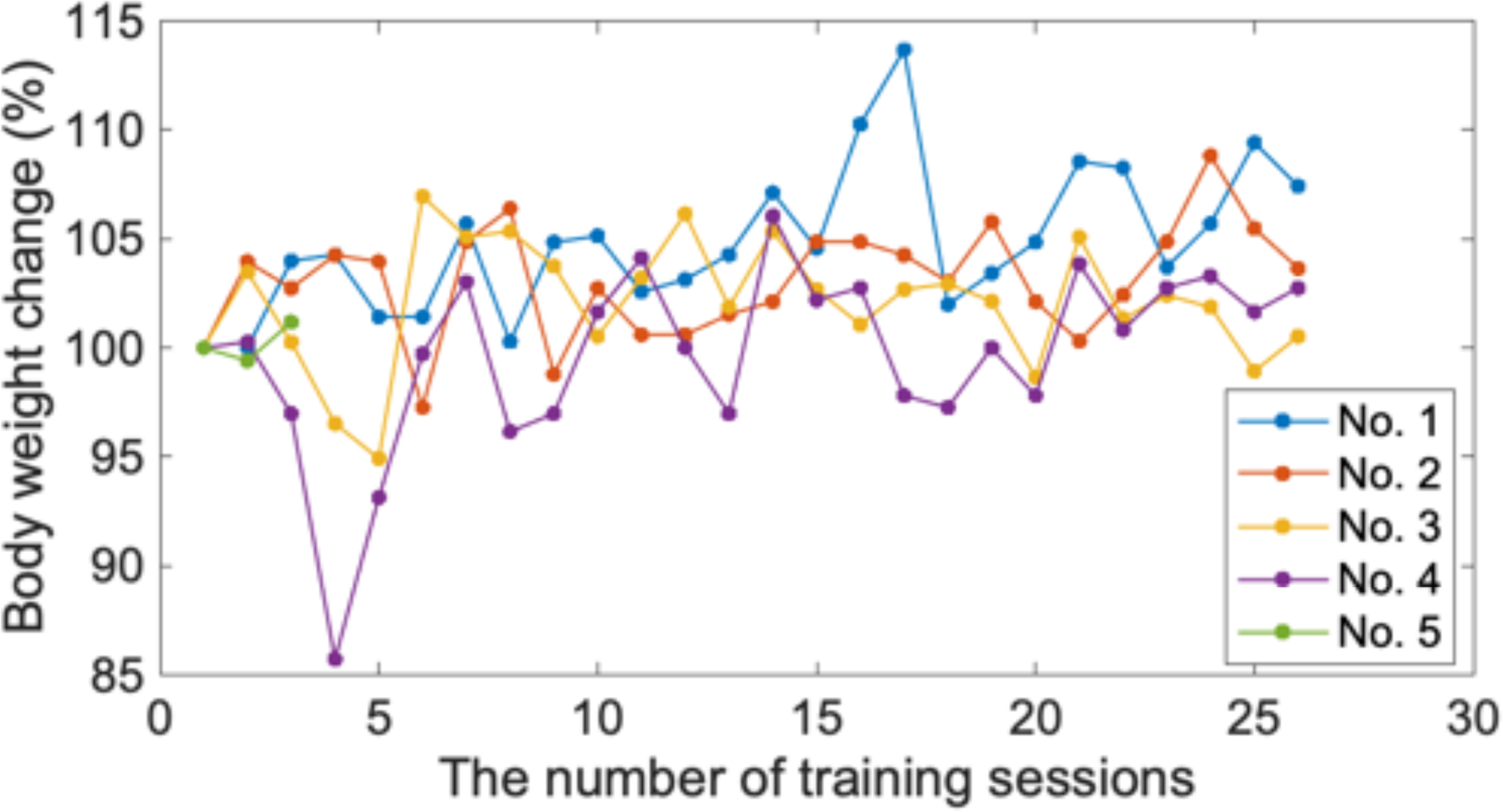
The body weight change of three animals during the acclimatization procedure. The acclimatization was conducted 26 times. Body weight was stable throughout the acclimatization.

In contrast, after acclimatization, the animals moved only 0.033 mm in the left-right direction, 0.068 mm in the vertical direction, and 0.107 mm in the front-back direction, indicating significantly decreased vertical movement (t-test, p < 0.01). In addition, careful observation via video recording confirmed that the marmoset was calm after acclimatization (Supplemental Video 2). The head movements of each animal during the fMRI scan are shown in Fig. 5. The animal movements were minimized using head restraints. The maximum estimated distance of reduced head translation among all marmosets was 0.32 mm for the x, y, and z directions, respectively. The maximum estimated degree of head rotation (pitch, roll, and yaw) was 0.94°.

**Figure 5.**
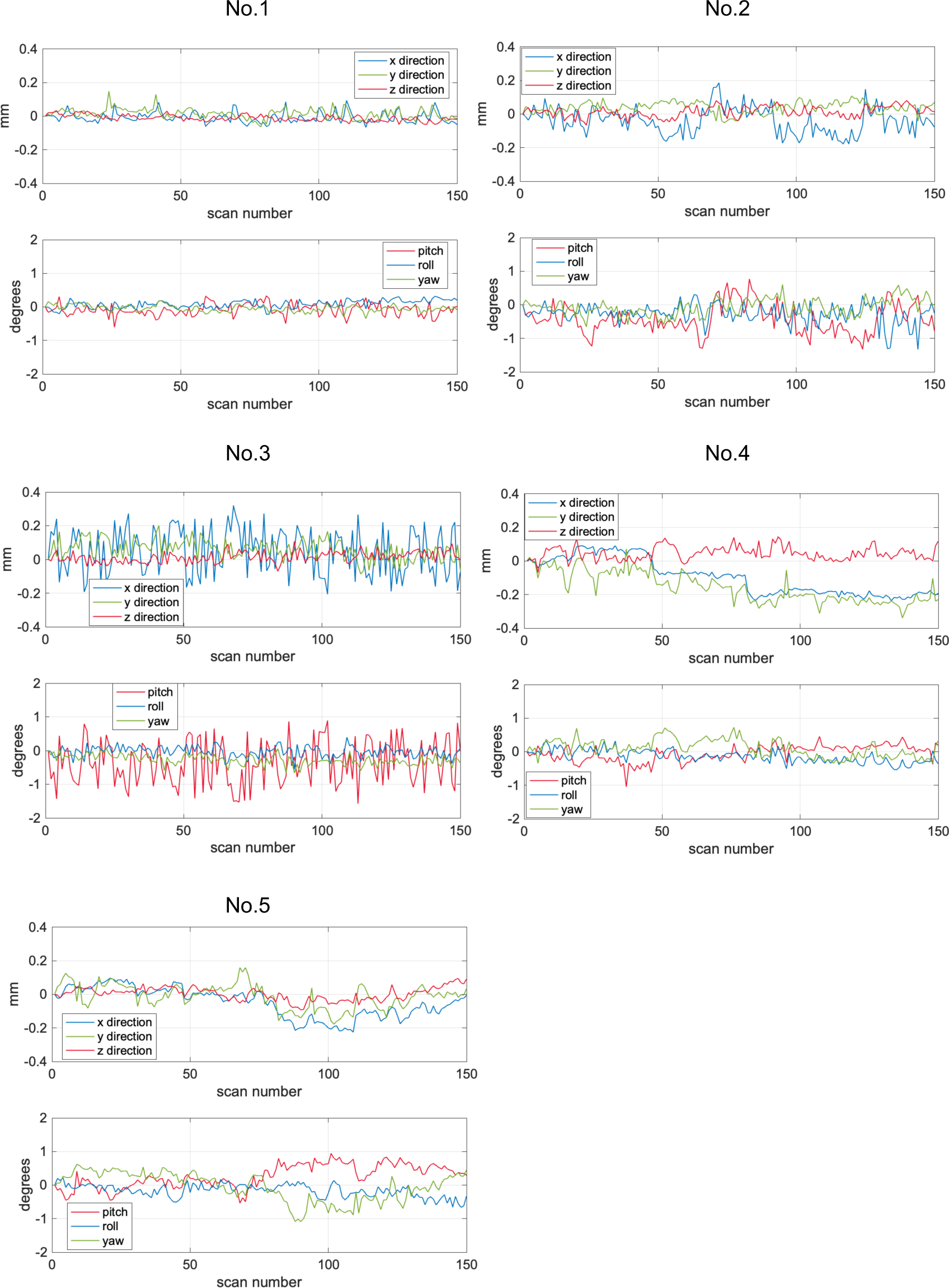
Estimates of head movement during the fMRI scans of five marmosets. Translation (top) and rotation (bottom) parameters are shown. The column indicates the number of the scan. The row indicates the translation (mm) above and the rotation (degrees) below.

### fMRI of auditory response

Fig. 6 shows that certain regions of the marmoset brain were significantly activated in response to each sound. When the marmosets heard the phee call, the regions included those in the auditory cortex, such as the primary auditory cortex (A1), auditory cortex middle lateral area (AuML) on the left side, auditory cortex anterolateral area (AuAL) on the left side, parts of the orbitofrontal cortex (A11/13), frontal pole (A10), and anterior cingulate (A24b). The prefrontal cortex (A11/13, A10, and A24b) was much more activated than the auditory cortex. In contrast, when marmosets heard the reverse sound of the phee call, fewer activation areas were observed than in normal phee calls, no significant activation in the prefrontal cortex, and slight activation in the auditory cortex rostral area (AuR) was observed.

**Figure 6.**
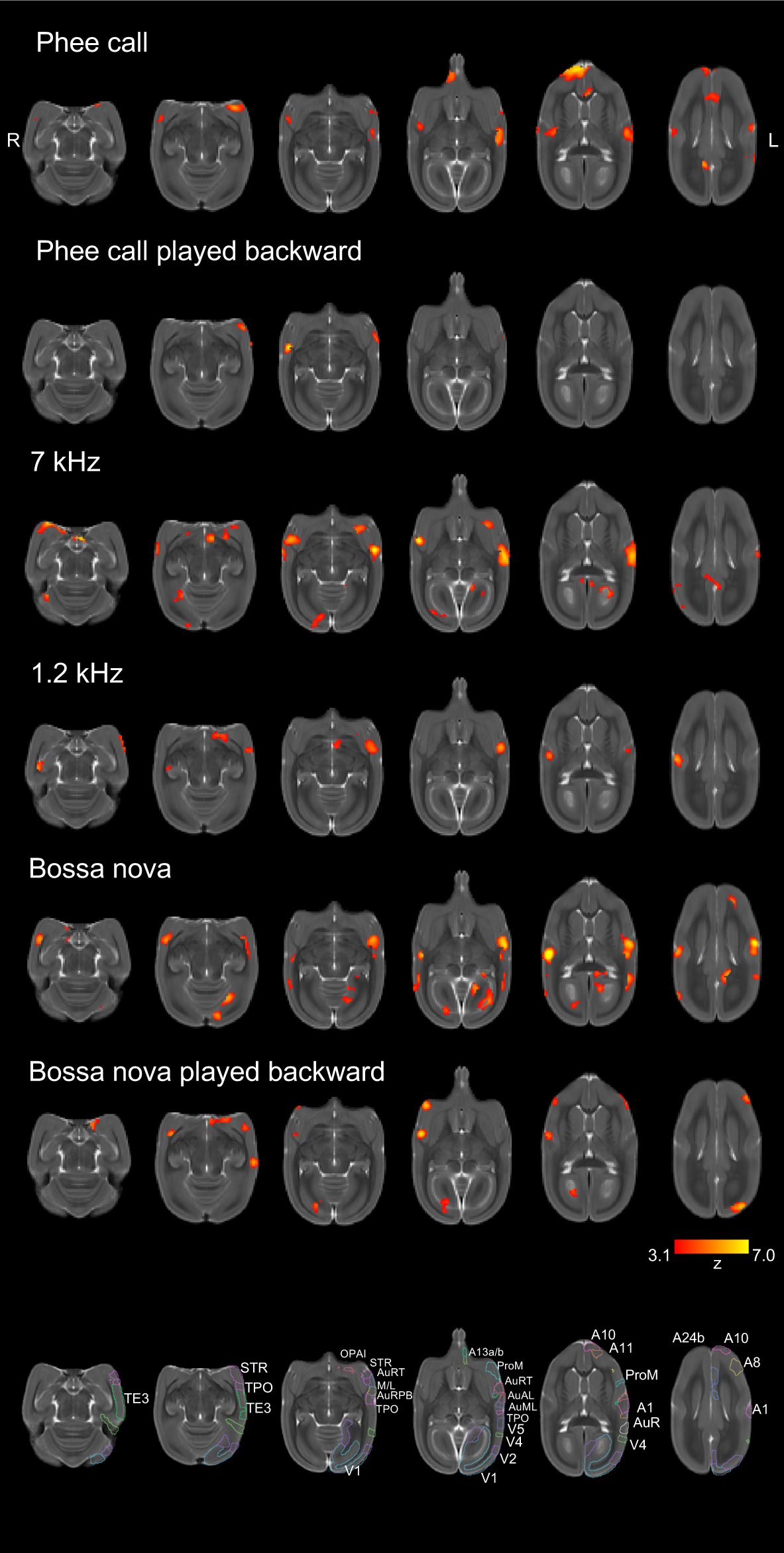
Cortical BOLD responses to auditory stimuli. The color map on axial slices of T2 weighted brain image shows the areas where statistically significant activation was observed for individual stimuli. From top to bottom: Phee call, Phee call played backward; 7 kHz, 1.2 kHz, Bossa nova, Bossa nova played backward. The bottom is a labeled T2 weighted image.

A10: Area 10 of the cortex; A8: Area 8 of the cortex; A1: Auditory cortex primary area; A24b: Area 24b of the cortex; ProM: Prosocortical motor region; AuR: Auditory cortex rostral area; V4: Visual area 4; A13b: Area 13 of the cortex; AuRT: Auditory cortex rostrotemporal part; AuAL: Auditory cortex anterolateral area; AuML: Auditory cortex middle lateral area; TPO: Temporo-parieto-occipital association area; V5: Visual area 5; V2: visual area 2; V1: Visual area 1; OPAI: Orbital periallocortex; STR: Superior temporal rostral area; AuRT, M/L: Auditory cortex rostrotemporal, medial/lateral area; AuRPB: Auditory cortex rostral parabelle; TE3: Temporal area 3.

When marmosets heard a 7 kHz pure tone, the AuAL, rostrotemporal lateral/medial area (AuRTL/AuRTM), and caudal and lateral parts of A1 were activated. The temporoparieto-occipital association area (TPO) and orbital periallocortex (OPAI) were also activated. Meanwhile, in 1.2 kHz tones, the bilateral A1, auditory cortex caudomedial area (AuCL), AuR, auditory cortex rostromedial area (AuRM), and auditory cortex rostrotemporal area (AuRT) were activated. The BOLD response was similar at 7 kHz and 1.2 kHz, as the TPO was also activated, particularly on the left side.

On hearing the bossa nova music, the bilateral auditory cortex, especially A1, the auditory cortex caudolateral area (AuCL), and the AuR on the left side were activated. In addition to the auditory cortex, the left dorsolateral prefrontal cortex (A8), inferior temporal cortex (TE3), primary visual cortex (V1), second visual cortex (V2), and fourth or fifth visual cortices (V4/V5) were also activated. Meanwhile, the regions activated by bossa nova being played backward were the auditory cortex, right side of A1, AuR, superior temporal rostral area (STR), V1/V2, and precentral opercular cortex (ProM). Compared to forward bossa nova music, the brain activation elicited by backward bossa nova was weak.

## Discussion

We developed a novel holder applicable to small-aperture MRI scanners to perform fMRI on awake marmosets non-invasively. The MRI was performed using an RF coil with an inner diameter of 72 mm, which provided less space to fix the head. We demonstrated that marmoset body and head movements were sufficiently minimized to conduct fMRI after acclimatization to the holder. The body weight change data suggested that this holder and the acclimatization were less stressful to the marmosets. Depending on the experimental design, it is possible to add stimulation devices to conduct fMRI with various types of stimulation using this holder.

Anesthesia or surgery was not necessary to perform fMRI with this holder. Therefore, the holder not only decreases the risk of anesthesia accidents but also decreases MRI artifacts. In general, skull-attached chambers are vulnerable to magnetic susceptibility image artifacts because of differences in magnetic susceptibility between the chambers, air, and adhesives (Schaeffer et al., 2020a). Additionally, the use of skull-attached chambers requires surgical intervention and competes with the advantages of non-invasive MRI. Our holder does not require surgery; thus, we do not need to accommodate such artifacts and can maximize the advantages of non-invasive MRI.

This holder does not require helmets for individuals. U-pins were used to fix the head and were resinous, so they bent slightly along the shape of the head. Using this feature and selecting the size of the U-pins on the head, we minimized head movement using the same method for all five marmosets. This feature indicates the high capacity of our holder to fit individual head shapes. We did not select marmosets that were likely to show good characteristics for fMRI, such as gentleness and calmness when fixed. It would be easy to increase the number of animals for scanning because our holder does not need to rely on the animals’ temperaments to conduct fMRI. Furthermore, the fact that the holder can scan all animals is useful when they cannot be selected because of specific conditions, such as genetically modified disease models or marmosets raised in a special environment.

Sensory-evoked functional magnetic resonance imaging fMRI demonstrated the holder’s utility. Among other sensory stimulations, auditory stimulation was the most appropriate choice to evaluate performance because auditory stimuli require minimal devices (i.e., earphones) and allow non-invasive and remote stimulation for animals. We observed increased activity in the auditory cortex and regions associated with auditory processing, which differed depending on the degree of ecological relevance.

A feature of acoustic processing with high ecological relevance (i.e., phee calls) is that input signals are processed to the auditory system via the frontal cortex (Jürgens, 2002). Our data showed that parts of the frontal cortex, such as the frontal pole (A10), which conveys the affective contents of the call (Reser et al., 2009), the orbitofrontal cortex (A11/13), and the anterior cingulate cortex (A24b), in addition to the auditory cortex, were activated, which was observed only for the phee call stimulus. This was consistent with the findings that activation of the prefrontal cortex, including the frontal pole and orbitofrontal cortex, increased in response to single phee calls using positron emission tomography (Kato et al., 2018), which was confirmed by the expression of the immediate early gene, c-fos (Miller et al., 2010a). In addition, activation of the anterior cingulate cortex for vocal perception has been confirmed using fMRI (Jafari et al., 2023) and functional ultrasound (Daniel et al., 2021). Using our holder, we acquired data sufficient to visualize the vocalization processing network consisting of the auditory cortex and frontal cortex using whole-brain fMRI.

In contrast, brain responses to low ecological relevance (pure tone) were observed only in the auditory cortex. Frequency-selective functional activation was observed in the auditory cortex. The response properties of neurons in marmosets are specific and depend on pitch, as identified in humans (Bendor and Wang, 2005). FMRI has confirmed that the caudal area in the auditory cortex is activated by high-frequency sounds (7 kHz. In contrast, the rostral area is activated by low-frequency sounds (1.2 kHz) (Toarmino et al., 2017). The results also show that the rostral area activated when marmosets heard low-frequency sounds. Without losing the frequency properties, our holder provided data that could differentiate frequency-selective functional activation to some extent.

Two different types of ecologically low-relevance sounds were evaluated to validate whether the marmosets in the holder showed a property-dependent brain response. The first is a sound that the marmosets have never heard (bossa nova). When marmosets heard bossa nova music, in addition to the auditory cortex, parts of the visual cortex (V1, V2, V4, and V5), the inferior temporal cortex (TE3), and the temporal-parietal-occipital association area (TPO) were activated. These regions are associated with visual processing but receive non-visual information from audition. It has been reported that when complex natural sounds are heard in daily life, such as singing birds and traffic noise, the early visual cortex is activated in addition to the auditory cortex (Lewis et al., 2004; Vetter et al., 2014).

Given that congenitally blind individuals also show such activation, neither visual experience nor visual imagery is required to develop this function (Vetter et al., 2020). Given the direct interconnections between the early visual cortex and the auditory areas, as observed in other primate cortices (Majka et al., 2018), the significant activation of the early visual cortex may be similar to the neural activity pattern in marmosets. The possible reason significant activation was observed in the visual cortex particularly when marmosets heard the bossa nova music would be that the neural response was particularly activated when they paid attention (Gandhi et al., 1999; Watanabe et al., 2011). Listening to music was the first experience for the marmosets, so it might attract marmosets more than the other two sounds and lead to activation of these regions.

The other sound was played after the phee call and bossa nova music. The backward playing of a phee calls has no social context. This sound and the 7 kHz sound had the same pitch as typical phee calls; however, neither induced significant frontal cortex activation, indicating that marmosets would not recognize the social context from these stimuli. Another feature of playing the sounds backward was the decline in whole-brain activity, which was observed in both phee calls and bossa nova. The previous study also reported that the marmoset primary auditory cortex (A1) responded more strongly to natural marmoset vocalizations than to reversed vocalizations (Wang et al., 1995). A human fMRI study also reported that the activation of brain regions elicited by backward-played sounds was lower than that by forward-played sounds (Lewis et al., 2004).

In summary, BOLD responses to each auditory stimulus captured their characteristics. The sound could reach the marmosets during experiments, and their specific brain activation was observed in response to the properties of each acoustic stimulus, as auditory cortical processing is property-dependent (King et al., 2018). We could infer that marmosets did not just recognize the pitch of the sound but also recognized the social context of the call despite the noise produced by the scanner. The holder was likely to function as intended without hindering vocalization-based stimuli. This demonstrates the high practicality of performing auditory-evoked MRI using our holder by sufficiently suppressing head motion.

Our study had some limitations. Our data did not replicate the previously reported brain activity. Activation of the dorsolateral prefrontal cortex(Kato et al., 2018) or ventrolateral prefrontal cortex (Miller et al., 2010a) in response to phee calls was not observed. In addition, although several activities in the auditory cortex were asymmetrical, significant activation of the inferior colliculus and medial geniculate nucleus, which comprise the auditory pathway, was not observed. The major reason for this is the insufficient spatial noise ratio in EPI. Because of size limitations, we used a commercially available volume coil rather than a phased array coil designed specifically for awake marmosets. Using phased-array coils greatly contributes to an increase in the spatiotemporal signal-to-noise ratio and resolution.

The activation of even small regions can be captured by optimizing the design to allow the application of a helmet-type coil for this bore size. Nevertheless, we observed significant activation of the prefrontal cortex, a critical area for perceiving calls in marmosets. In addition, when a phased-array coil designed for awake marmosets is unavailable, our data demonstrates that examining brain activity in response to auditory stimuli is possible.

We fulfilled our aim of developing a new holder for awake marmoset fMRI. We acquired reliable data from the brain activation of auditory responses and confirmed the usefulness of the holder. Given that the individual conditions of MRI specifications differ among institutions, the proposed holder would be flexible enough to adapt to the different conditions because the components of the holder are minimal and simple. This non-invasive holder has advantages in the long-term evaluation of model marmosets using genetic modification technologies and neurodegenerative disease models, including Parkinson’s disease and Alzheimer’s disease, to characterize changes in brain networks and understand the brain-gut relationship in nonhuman primates (Inoue et al., 2021; Sasaguri et al., 2022; Sato et al., 2016; Sato et al., 2020).

## CRediT authorship contribution statement

Fumiko Seki: Data curation, Formal analysis, Funding acquisition, Investigation, Project administration, Software, Validation, Writing – original draft. Terumi Yurimoto: Conceptualization, Funding acquisition, Investigation, Methodology, Resources, Software, Visualization, Writing – original draft. Michiko Kamioka: Data curation, Investigation, Visualization, Writing – review & editing. Takashi Inoue: Investigation, Supervision, Validation, Writing – review & editing. Yuji Komaki: Formal analysis, Resources, Visualization, Writing – review & editing. Atsushi Iriki: Investigation, Methodology, Supervision, Validation, Writing – review & editing. Erika Sasaki: Conceptualization, Funding acquisition, Methodology, Project administration, Resources, Supervision, Writing – review & editing. Yumiko Yamazaki: Conceptualization, Investigation, Methodology, Project administration, Supervision, Writing – review & editing.

## Data availability statement

The data supporting this study’s findings are available on request from the corresponding author. The parts of the holder exported as STereoLithography file format are available in Supplemental Data 1.

## Ethics approval statement

The Institutional Animal Care and Use Committee of the CIEA (approval numbers: 20068A, 21129A) approved all animal experiments.

## Patient consent statement

This study did not use human data.

## Abbreviations

fMRI: Functional magnetic resonance imaging
CIEA: Central Institute for Experimental Animals
CAD: computer-aided
EPI: echo-planar imaging
BOLD: blood oxygenation level-dependent
TR: repetition time
TE: echo time
FOV: field of view
T2WI: T2-weighted images
RARE: rapid acquisition with relaxation enhancement
A1: primary auditory cortex
AuML: auditory cortex middle lateral area
AuAL: auditory cortex anterolateral area
AuR: auditory cortex rostral area
AuRTL/AuRTM: auditory cortex rostrotemporal lateral/medial area
AuCL: auditory cortex caudomedial area
AuRM: auditory cortex rostromedial area
AuRT: auditory cortex rostrotemporal area
STR: superior temporal rostral area
ProM: precentral opercular cortex
TPO: temporal-parieto-occipital association area

## Acknowledgments

This research was supported by the Brain Mapping by Integrated Neurotechnologies for Disease Studies (Brain/MINDS), “Study of developing neurodegenerative model marmosets and establishment of novel reproductive methodology (JP19dm0207065)” and “Platform for marmoset research support” (JP19dm0207068) from the Japan Agency for Medical Research and Development (AMED) and Grant-in-Aid for Scientific Research A “Development of an Automated Behavioral Analysis System for Evaluation of Genetically Modified Disease Model Marmosets” JSPS KAKENHI Grant Number JP21H04756 to ES; Grant-in-Aid for Early-Career Scientists “Effects of transient alcohol exposure to the fetus in early pregnancy on brain development”; JSPS KAKENHI Grant Number JP20K16908 to TY; Grant-in-Aid for Early-Career Scientists “Unveiling vulnerability of the neural connectivity controlling nicotine intake in marmoset brain development”; JSPS KAKENHI Grant Number JP19K16031 to FS.

---

Supplementary Figure 1. Positions of earphones to provide auditory stimuli using a mock monkey. (A) Top view and (B) side view.

Supplementary Figure 2. Position of earphones for acclimatization of fMRI sounds in a mock monkey. (A) Top view and (B) side view.

Supplementary Figure 3. Rendered CAD drawing of the animal holder.

Supplementary Video 1. Demonstration of the marmoset holder using a mock monkey.

Supplementary Video 2. Observation before and after the acclimatization to the holder. A. Before acclimatization, the eyes were closed soon after the marmoset was placed in the holder. The animal did not show any interest in feeding. B. After acclimatization, the animal appeared slightly sleepy but awake. He licked when fed.

Supplementary Data 1. The individual parts of the holder are exported in STereoLithography file format.

