## Supplementary figures and images for "Development of a non-invasive novel individual marmoset holder for evaluation by awake functional magnetic resonance brain imaging"

### Supplemenetal figure 1

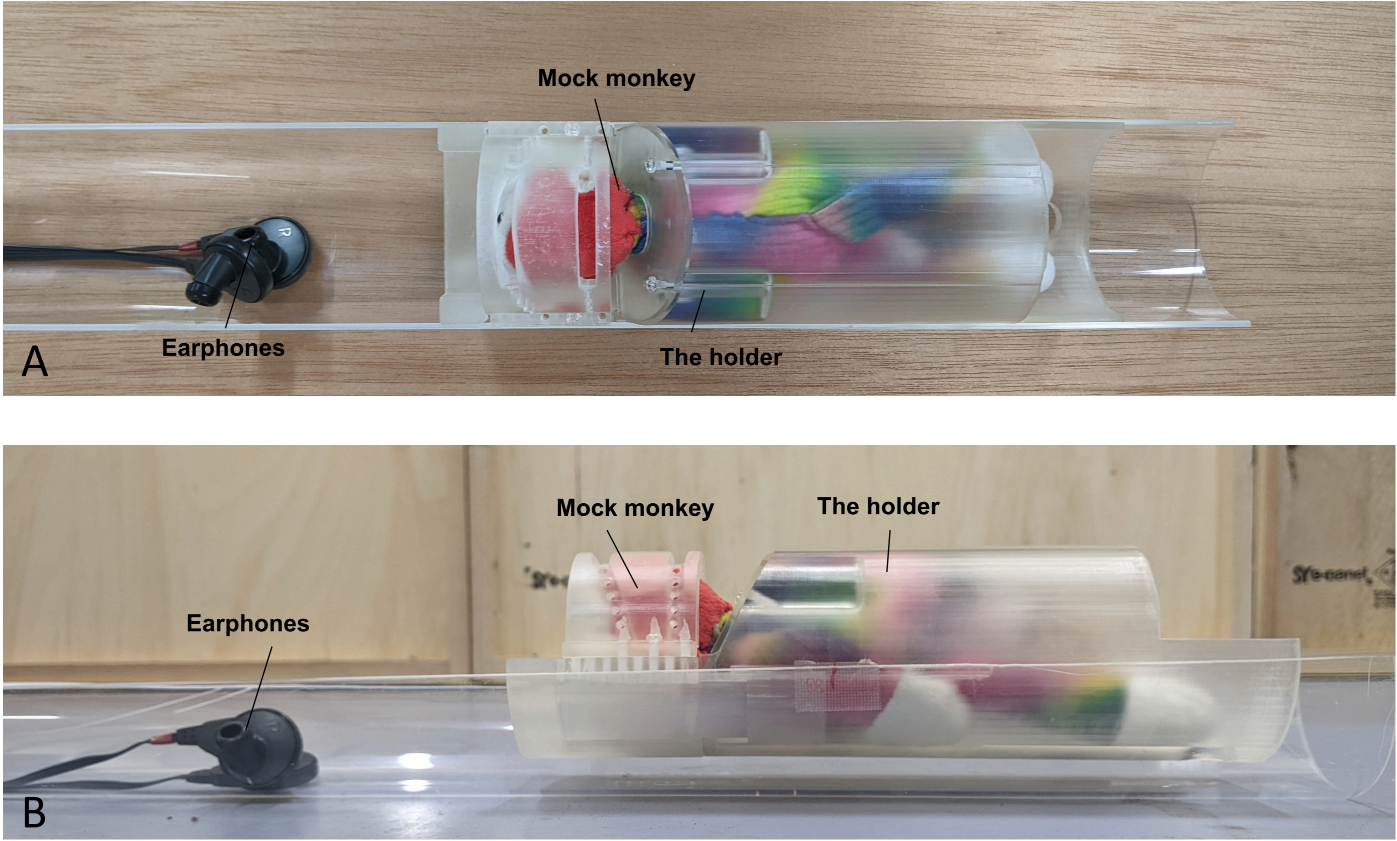

### Supplemenetal figure 2

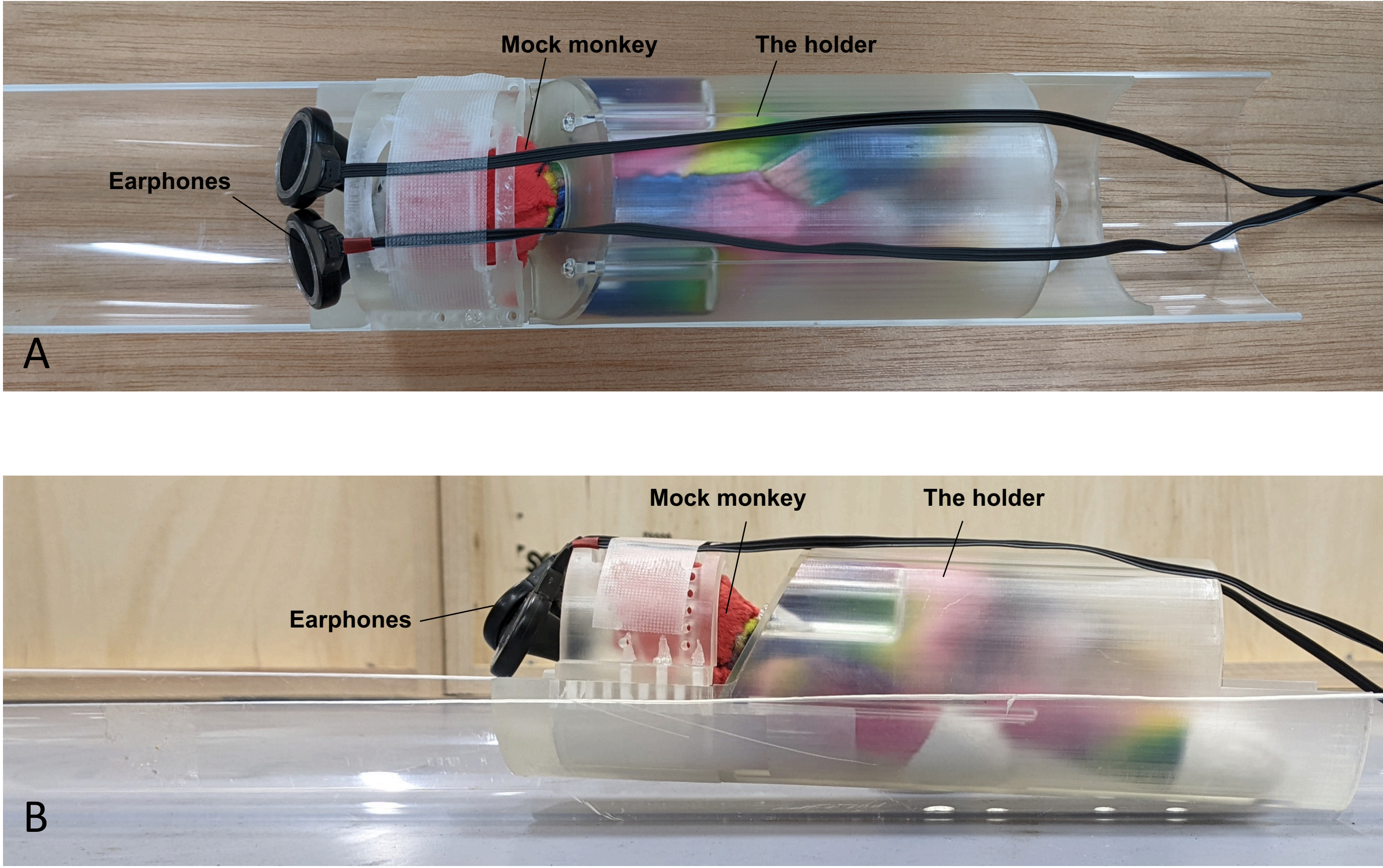

### Supplemenetal figure 3

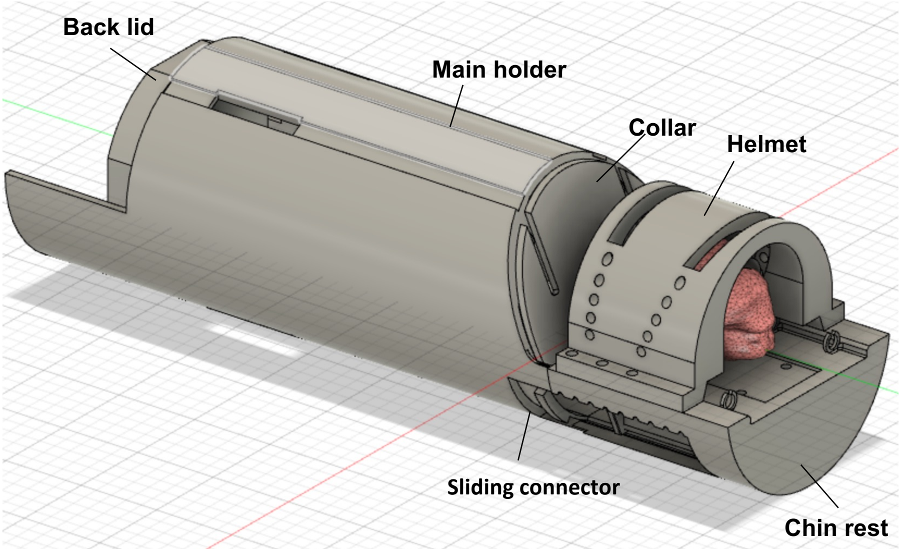
